# Developing a 670k genotyping array to tag ∼2M SNPs across 24 horse breeds

**DOI:** 10.1101/112979

**Authors:** Robert J. Schaefer, Mikkel Schubert, Ernest Bailey, Danika L. Bannasch, Eric Barrey, Gila Kahila Bar-Gal, Gottfried Brem, Samantha A. Brooks, Ottmar Distl, Ruedi Fries, Carrie J. Finno, Vinzenz Gerber, Bianca Haase, Vidhya Jagannathan, Ted Kalbfleisch, Tosso Leeb, Gabriella Lindgren, Maria Susana Lopes, Nuria Mach, Artur da Câmara Machado, James N. MacLeod, Annette McCoy, Julia Metzger, Cecilia Penedo, Sagi Polani, Stefan Rieder, Imke Tammen, Jens Tetens, Georg Thaller, Andrea Verini-Supplizi, Claire M. Wade, Barbara Wallner, Ludovic Orlando, James R. Mickelson, Molly E. McCue

## Abstract

**Background:** To date, genome-scale analyses in the domestic horse have been limited by suboptimal single nucleotide polymorphism (SNP) density and uneven genomic coverage of the current SNP genotyping arrays. The recent availability of whole genome sequences has created the opportunity to develop a next generation, high-density equine SNP array.

**Results:** Using whole genome sequence from 153 individuals representing 24 distinct breeds collated by the equine genomics community, we cataloged over 23 million *de novo* discovered genetic variants. Leveraging genotype data from individuals with both whole genome sequence, and genotypes from lower-density, legacy SNP arrays, a subset of ∼5 million high-quality, high-density array candidate SNPs were selected based on breed representation and uniform spacing across the genome. Considering probe design recommendations from a commercial vendor (Affymetrix, now Thermo Fisher Scientific) a set of ∼2 million SNPs were selected for a next-generation high-density SNP chip (MNEc2M). Genotype data were generated using the MNEc2M array from a cohort of 332 horses from 20 breeds and a lower-density array, consisting of ∼670 thousand SNPs (MNEc670k), was designed for genotype imputation.

**Conclusions:** Here, we document the steps taken to design both the MNEc2M and MNEc670k arrays, report genomic and technical properties of these genotyping platforms, and demonstrate the imputation capabilities of these tools for the domestic horse.

## Background

Soon after the horse reference genome from Twilight, a female Thoroughbred, was sequenced using Sanger technology [1], a genotyping array (Illumina EquineSNP50 BeadChip) was developed to enable whole genome mapping using ∼50k (54,602) single nucleotide polymorphism (SNP) markers from low-coverage Sanger sequence of 7 horses representing 7 different breeds (an Andalusian, Arabian, Akhal-Teke, Icelandic, Standardbred, Thoroughbred and a Quarter Horse) [2]. Shortly thereafter, a slightly higher density array (Illumina Equine SNP70 BeadChip) with ∼65k (65,157) informative SNP markers was developed. These widely used, now legacy, SNP arrays (with a combined 74,056 unique SNPs, ∼74k hereafter [3]), have successfully enabled genetic studies examining domestication and selection [4,5], disease and performance trait mapping [6–15], and population structure and dynamics [2,16,17]. However, extensive population structure and the low extent of linkage disequilibrium (LD) existing in many horse breeds severely limits conventional mapping approaches with the relatively low SNP density on the current SNP array [17]. Since the initial Sanger shotgun sequencing of Twilight, whole genome sequence (WGS) has been generated for hundreds of horses [15,18–25], prompting the development of a new higher-density genotyping array for the horse.

Here we describe the steps taken, including careful and extensive variant filtering, to create both a 2 million (2M) SNP array and a 670 thousand (670k) SNP array from over 23 million variants discovered from whole genome sequence of 153 horses representing 24 breeds (See Table S1). Genotypes from the 2 million SNP genotyping array (MNEc2M) in a cohort of 332 horses, from 20 actively-researched or economically-important breeds that represent known genetic diversity in the domestic horse [17], were used to select SNPs for inclusion on the commercially-available 670k SNP array (MNEc670k). The MNEc670k array was designed for accurate genotype imputation up to the higher density, 2M SNP set present on the MNEc2M array. We report summary statistics, broken down by breed, for both arrays as well as preliminary results on genotype imputation performance from the MNEc670k array to the SNP density on the MNEc2M array.

## Results

### Variant discovery from whole genome sequence

Whole genome, 100 base pair (bp), paired-end Illumina HiSeq reads were generated for 153 horses (including Twilight), representing 24 distinct breeds, at a depth between 1.7X and 64X, with a median depth of 13X (Table S1). Read mapping was performed using the PALEOMIX genome mapping protocol [26] to efficiently process samples in parallel and to assess individual sample quality. Each sample was mapped to the EquCab2.0 reference genome, which had been extended with an additional 7,850 *de novo* assembled scaffolds (See Methods), to produce a total of nearly 48 billion unique reads aligned to the nuclear genome (See Table S1). Variants were identified by extending the PALEOMIX framework to identify SNPs using two variant callers (see Methods). To maximize efficiency of variant calling in individual breeds, and to minimize bias due to variable sequencing depth of coverage, individuals were broken up into 16 variant calling groups (see Table S1, Variant Calling Group columns) by estimated depth of coverage and breed. Variants were called using permissive parameters in both the GATK UnifiedGenotyper [27] as well as SAMtools ‘mpileup’ utilities [28]. Approximately 23 million potential SNPs were in the intersection of SNP sets called by GATK and SAMtools. These ∼23 million SNPs were kept for further analysis and validation (See Table 1; Supp. File 1).

### Precision of GATK QUAL scores for variants identified by both callers

We evaluated the performance of GATK QUAL scores by comparing genotypes generated from WGS to genotypes generated from the legacy Equine 54K SNP chip in 23 horses at two different sequencing depths. Variants detected on chromosome 1 (ECA1) by both WGS and 54k SNP chip were ranked by decreasing QUAL score. The proportion of genotype calls that agreed between 54K and WGS (i.e., precision) was compared based on ranked QUAL score (i.e., recall) [29]. For each ranked variant we evaluated the precision at that point, which is the cumulative proportion of genotypes that agree between the two genotyping methods (Fig. 1). We note that this approach evaluates the concordance between the two genotyping technologies and is thus unable to assess the accuracies of the underlying genotypes themselves. However, this approach does take into account the imbalance of false positives/negatives between genotypes called by WGS and by the 54K SNP chip [30].

**Fig. 1.**
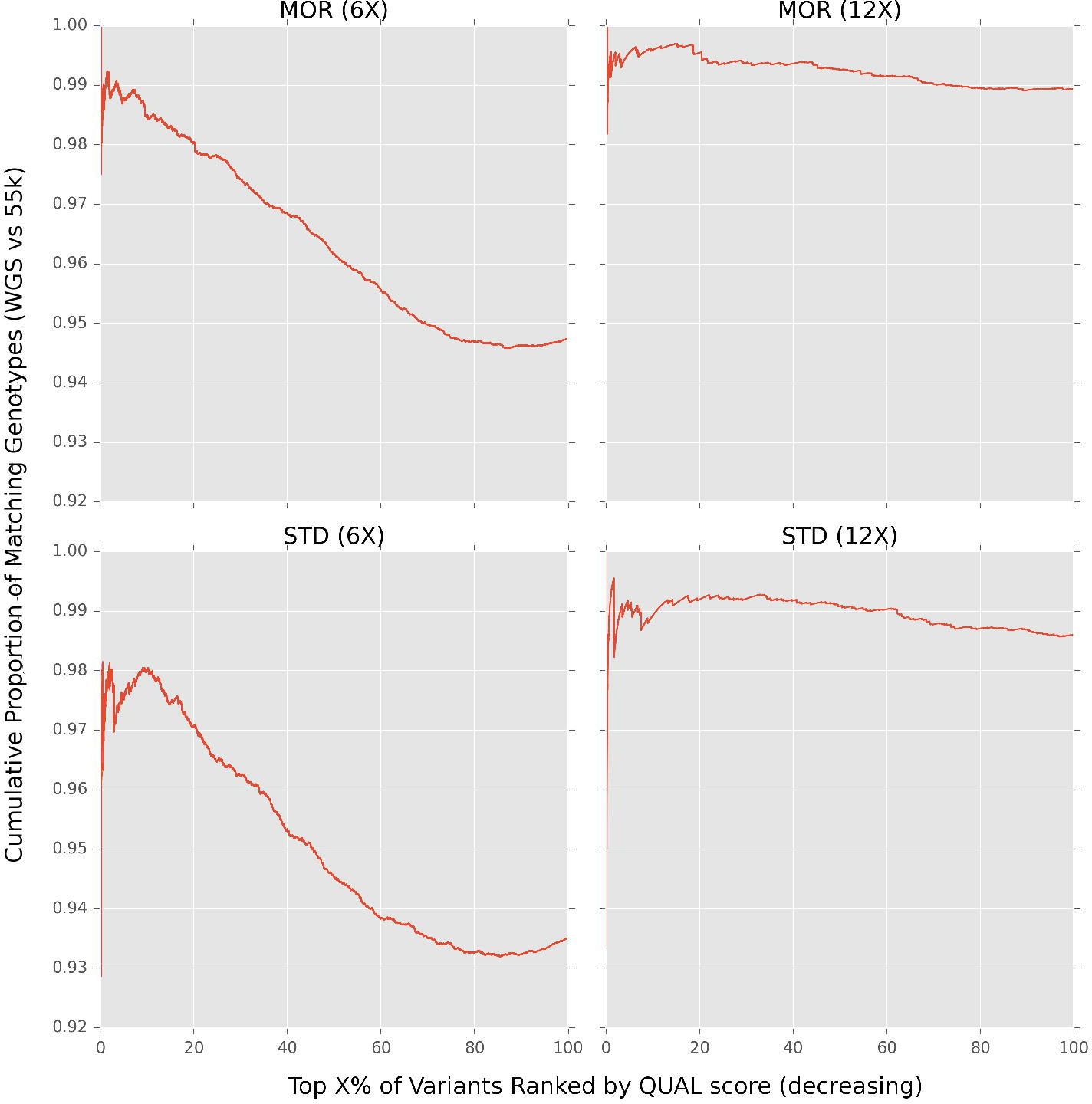
*Comparing high QUAL genotypes called de-novo to the SNP50 array in Morgans and Standardbreds*. SNPs on chromosome (ECA1) 1 with genotype calls from both WGS and SNP50 array were ranked by decreasing QUAL score for Morgans (MOR) at 6X (3,759 SNPs; 12 individuals) and 12X (3,751 SNPs; 6 individuals) coverage, and Standardbreds (STD) at 6X (4,422 SNPs; 5 individuals) and 12X (4,380 SNPs; 3 individuals) coverage. The cumulative proportion of genotypes that agree between platforms was compared based on ranking *de-novo* variants by QUAL score. Variants with missing data on either platform were excluded.

Considering 100 percent of variants detected by both technologies, genotypes had an overall precision of ∼99% for 12X-called variants and 94-95% for 6X-called variants (See Fig. 1, x-axis). This is consistent with results from a similar study that compared *de novo* variant genotypes to array based genotypes in the Franches-Montagnes horse breed [25]. Yet, many SNPs with very high QUAL scores (e.g., SNPs within the top 10 percent; Fig. 1) had disagreeing genotype calls between WGS and 54K SNP chip. Additionally, when considering between 80-100% of variants ranked by QUAL score (Fig. 1), in both 6X MOR and STD comparisons, the proportion of matching genotypes increases, indicating that there are many variants with low QUAL scores and high precision. These results indicated that QUAL scores alone did not adequately rank variants, and that additional metrics were necessary to improve the reliability of SNPs ultimately chosen for the higher density genotyping arrays.

### Identifying gold standard reference set for variant recalibration

In addition to QUAL type quality scores, GATK outputs additional metrics such as depth, quality of depth, Fisher strand position, map quality rank sum, and read position rank sum for each variant based on read statistics (See Methods for details of these metrics). These values were used to train linear mixed models, using the GATK VariantRecalibrator [31], to assign a composite quality score (VQSLOD) to help detect type I and II errors.

Training these models required a “gold standard” reference set of known genotypes across multiple individuals. Lacking this resource in the domestic horse, we defined three high confidence, putative “gold standard” datasets with which to train the VariantRecalibrator: 1) SNPs on the legacy SNP50 chip; 2) WGS variants which were seen in four or more (4+) calling groups (Table S1); and 3) WGS variants that were in the top one percent of QUAL scores. Models were trained on features from “gold standard” variants present on chromosomes 2-32 and used to calculate VQSLOD scores for SNPs on chromosome 1. To assess the performance of VSQLOD scores generated by each training set, scores from each group were compared to each other in addition to QUAL scores using horses genotyped on the 54K SNP Chip.

SNPs on chromosome 1 (ECA1) were ranked either by decreasing QUAL or by decreasing VQSLOD score generated from each of the three gold standard training groups. Fig. 2 shows the proportion of matching genotypes between WGS and SNP50 platforms for each of the four groups. With the exception of 6X Morgans, SNPs with high VQSLOD scores agreed more often with genotypes called on the 54K SNP chip compared to SNPs with high QUAL scores alone. For example, the top 10% highest scoring VQSLOD scores had a higher concordance rate than QUAL scores in all cases except for MOR6X, where only two of the recalibrated scores marginally outperformed QUAL scores (See Fig. 2; see discussion). Additionally, VQSLOD variants did not drop below the overall discordance rate, showing an overall better ranking than QUAL scores alone (red curve, Fig. 2).

**Fig. 2.**
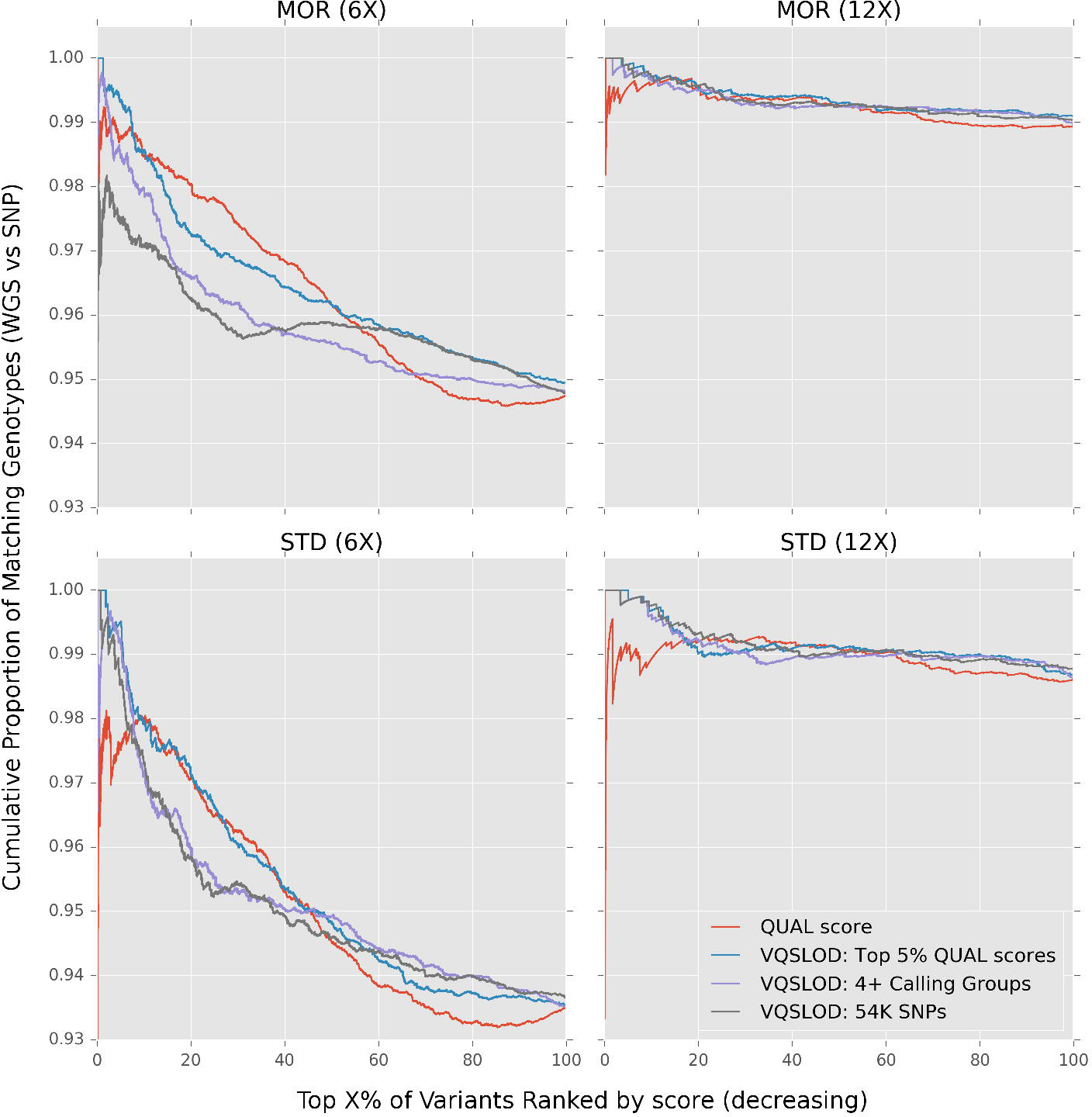
*Comparing QUAL ranked SNPs to VQSLOD ranked SNPs*. VQSLOD scores were calculated from three different “gold standard” reference groups in Morgans (MOR) and Standardbreds (STD) using GATK VariantRecalibrator. Compared to QUAL scores (red line), high VQSLOD scored variants (top 10% variants by score) have a lower number of mismatched genotypes across the SNP50 BeadChip and variants discovered *de novo*.

While an exact differentiation between the gold-standard training groups remains unclear (See Discussion), recalibrated VQSLOD scores were better able to differentiate high scoring, false positives than QUAL scores alone, making them a more informative metric for selecting the final SNPs for use on the high-density commercial array. VQSLOD scores were calculated for all 23 million variants, in all remaining horses, using WGS variants called in the 4+ breed calling groups (described above) as the training set (candidate 23M SNP set). Scored variants were then filtered in several steps (described below) to select the subset of SNPs to be included on the new higher density arrays.

### Preliminary 5M SNP selection for genotyping probe design

We analyzed the 23M candidate SNP set with the goal of generating a target set of ∼5 million high-confidence SNPs that would be compatible with array design. Following criteria provided by Affymetrix (now Thermo Fisher Scientific), 780 SNPs were filtered because they fell within repetitive regions; 9,342,733 SNPs were filtered because they were within 20 base pairs (bp) of another variant; and 4,858,273 SNPs were filtered for having a minor allele frequency below 1% in the 153 WGS horses. After filtering, approximately 10 million SNPs remained for further design selection (hereafter referred to as the 10M SNP Set, Table 1). Of these ∼10 million array-compatible candidates, 48,485 SNPs (65% of the total) overlapped with the sites available from the ∼74k legacy SNP sets (although all legacy SNP sites were included in MNEc2M/670k array design). Supp. Fig. 1 shows the alternative allele frequency distribution of the 23M *de novo* and 10 million design candidate SNPs (10M SNP set). The mean allele frequency is below 0.1, though there is a long tail of SNPs with a high alternative allele frequency. Interestingly, there are 445,421 SNPs (23M set) and 46,455 SNPs (10M set) where the alternative allele frequency is 1.0. This is likely due to errors in the EquCab 2.0 reference sequence (Sanger reads from Twilight) as Twilight’s whole genome Illumina sequence data (collectively at more than 25X), and whole genome sequences of all other horses, contained the alternate allele. These SNPs were excluded from array design.

To ensure backwards compatibility, we started with the ∼74k SNPs that were present on the two legacy (54K/65K) arrays. To fill the remaining target of ∼5 million probe design candidates, we added SNPs from the 10M SNP set based on even genome distribution, informativeness among the variant calling groups (See Table S1), and linkage disequilibrium (LD). LD was calculated for all pairs of SNPs within 10kb of each other throughout the genome. SNPs that were in high LD (r^2^ > 0.90) were filtered based on whether or not they were present in draft or pony variant calling groups to control for the fewer number of samples used in variant discovery. Previous SNP designs [2] show an underrepresentation of informative SNPs from these groups, which leads to poor mapping resolution in draft and pony breed groups (See Table S2). SNPs discovered in both draft and ponies were prioritized over those discovered in one group and over those absent in both. If SNPs were discovered in an equal number of priority groups, VQSLOD scores were used to break ties.

After applying these SNP candidate criteria, 5,443,950 SNPs (5M set, see Table 1) were kept, of which 2,199,467 SNPs occurred in pony calling groups and 2,782,917 SNPs occurred in draft calling groups. A total of 1,695,347 SNPs occurred in both ponies and drafts. Flanking sequences (35 bp upstream and downstream) for these 5M filtered SNPs were submitted to Affymetrix for SNP probe design analysis.

### SNP selection for the high density MNEc2M SNP array

SNP conversion recommendations for the 5M SNPs were provided from Affymetrix in both forward and reverse strand directions. SNPs were assigned to one of four groups based on decreasing probability of successful probe design: ‘recommended’, ‘neutral’, ‘not recommended’, and ‘not possible’. To achieve an even distribution of ∼2 million SNPs, the equine reference genome (∼2.7 Gb) was divided into approximately 54,000 50 kb windows and a target of 37 SNPs per window was established. SNPs within each window were chosen for inclusion in the MNEc2M SNP set using a greedy algorithm. Briefly, the ∼74,000 SNPs on previous generation arrays, as well as SNPs within the equine major histocompatibility complex (MHC) region (ECA20:28.7-33.6Mb), were given VIP status and were automatically included in both forward and reverse strand directions. SNPs were added to windows until the target of 37 SNPs was met. If a window had more than 37 candidates, SNPs were selected based on by Affymetrix recommendation group (See Methods for details). In total, given the above criteria, 2,001,826 high quality SNPs (2M SNP set) were chosen and submitted for probe design to comprise the MNEc2M genotyping array (Supp. File 2).

### SNP selection for the MNEc670k array

A cohort of 347 horses were genotyped on the MNEc2M high-density array using DNA isolated from blood (n=286) or hair roots (n=61) (see Methods). The ∼2M SNP genotypes were split across three different physical arrays. Genotypes were called using Affymetrix Power Tools and sample quality control was assessed according to Affymetrix best practices (See Methods). For each sample, the three physical arrays were assessed separately for quality control. In total, 320 samples passed in all three arrays while 27 samples had one or more arrays that did not pass quality control. Of the 27 samples that failed, 7 samples had two passing arrays, 5 samples had one passing array, and 15 DNA samples failed to meet genotyping quality control metrics on all three physical arrays. Failed arrays were removed from the analysis. If a sample had at least one array, genotypes from those arrays were retained. In total, viable genotypes were produced for 332 horses.

Of the failed arrays, 25 had DNA isolated from hair roots and 2 had DNA isolated from blood. Hair root DNA had a lower average DNA concentration (Pico Green) when re-hydrated (2.7 ± 2.9 ng/µL) than did DNA samples isolated from blood (43.1 ± 55.5 ng/µL). To determine if sample origin (blood versus hair roots) or DNA concentration, or both, were associated with failure to pass genotyping quality control metrics, a logistic regression for sample success on DNA concentration and blood/hair status was performed. (Supp. Fig. 6). Both DNA source (p ≤ 4.05e-04) and DNA concentration (p ≤ 2.77e-10) significantly influenced the probability of samples producing quality genotypes. Both factors also had substantial coefficients in the model indicating a large magnitude of effect. In the logistic regression, the factor indicating hair root had a strong negative model coefficient of -2.70 while DNA concentration had a positive coefficient of 1.71. (see Discussion).

After quality control steps for samples, genotypes were called for all 2,001,826 SNPs. Genotypes were assessed for clustering quality using the *Metrics.R* script provided by Affymetrix (see Methods). In total, 92.2% of SNPs on the MNEc2M array passed quality control producing a set of 1,846,988 high-quality SNPs (1.8M; see Table 1) genotyped on the 332 horses remaining in the analysis (see Methods).

SNPs exclusive to a single WGS variant calling group were validated (polymorphic), on average, at a rate of 80%. Yet, validation rates were much higher in SNPs discovered in multiple calling groups (>96%) (See Table S3). Genotypes produced from the 332 horses with passing genotypes on the MNEc2M array (“SNP” genotypes) were combined with genotypes discovered from the 153 whole genome sequence horses (“WGS” genotypes) to create a dataset containing 1.8M genotypes for 485 horses (“WGS+SNP” genotypes). These variants were analyzed to select a subset of SNPs for inclusion on the MNEc670k array, with the intent that this array would be designed for genotype imputation to the full 2M SNP set.

To maximize the information content of the MNEc670k array SNPs, multi-marker r^2^ statistics were calculated on the 1.8M high-quality candidate SNPs to identify ‘tagging SNPs’ that allowed for efficient reconstruction of genomic haplotypes in the 485 horses both within and across breeds. Tagging SNPs that reconstructed haplotypes across all 485 horses (inter-breed tag SNPs) were identified using the software FastTagger [32]. In total, 355,903 tagging SNPs were needed to reconstruct haplotypes present across all the horses in the cohort with a minor allele frequency (MAF) > 0.01 and tag-SNP r^2^ > 0.99. These SNPs were included on the MNEc670k imputation array (see Table S4; Inclusion criteria: Inter).

Haplotype tagging SNPs were also examined at the breed-specific level. Horses were split into 15 *tagging breed groups* based on the minimal sample size necessary to perform SNP tagging (see Table S5). Tagging SNPs were identified separately in each tagging breed group using FastTagger, then a subset of population specific tag SNPs were identified using the software Multi-Pop-TagSelect [33]. Combined, a total of 1,754,075 SNPs were needed to reconstruct fine-level, breed-specific haplotypes in all of these breed groups (MAF > 0.10 and tag SNP r^2^ > 0.90; See Supp. File 3). Breeds varied in the number of tag SNPs needed to reconstruct haplotypes. Table 2 shows the number of tagging SNPs required to reconstruct haplotypes in each of the tagging breed groups. The Ponies, Draft and Quarter Horses (tagging breed groups) required the most tagging SNPs, each requiring over 350,000 SNPs to reconstruct breed-specific haplotypes, while Thoroughbred, Icelandic and Lusitano tagging breed groups each required less than 150,000 tag SNPs, to reconstruct haplotypes.

Tagging SNPs that were informative in 5 or more tagging breed groups were included on the MNEc670k array (n=206,822; see Table S4; inclusion criteria: Intra). An additional 13,993 SNPs that tagged haplotypes in four of the breeds requiring a larger number of tag SNPs (Quarter Horse, Pony, Morgan, Standardbred) were also included in the array (inclusion criteria: Diverse). Additionally, 7,394 SNPs were included due to their location within the equine MHC region (inclusion criteria: MHC), 16,398 SNPs were included to increase SNP density in 12,104 (24.3%) 50 kb genomic windows to at least 8 SNPs (inclusion criteria: Density), and 70,295 SNPs were retained for backwards compatibility with legacy arrays (inclusion group: VIP). Collectively, 670,805 SNPs were included on the MNEc670k commercial array.

### Imputation accuracy from the MNEc670k SNP set to the MNEc2M SNP

Genotype imputation accuracy from the MNEc670k array to the MNEc2M array was quantified using the 485 horse reference population. For each of the tagging breed groups a subset of 1/3 random individuals were masked down to the MNEc670k SNP set. Genotypes were then imputed to the MNEc2M SNP using the reference population after removing the individuals being imputed (See Methods). Imputation accuracy was measured as total genotype concordance across imputed individuals (correctly inferred genotypes/total number of imputed genotypes; see Methods for details).

Genotype imputation accuracy from the MNEc670k SNP set to the MNEc2M SNP set ranged between 96.6 and 99.4% in the 15 breeds tested (see Table 3). Tagging breed groups with over 99% mean imputation accuracy included Arabians, Belgians, Lusitano, Maremanno, Pony and Thoroughbred. No tagging groups were below 98% with the exception of the Draft group. This breed group contained both continental European draft breeds as well as British Isles draft breeds which have been previously shown to have distinct sub-population structure[17]. Random sampling for imputation validation in the Draft group included the only two Percheron samples (M1542 and M1545), which underperformed (average 92.6% accuracy) compared to the other Draft samples in the imputed group (average 98.5% accuracy), performing similarly to the other heavy horse breeds (e.g. Belgian). We suspect that an increased representation of Percheron samples in the reference population would increase imputation performance for individuals within that tagging breed group (See Discussion).

The effect of allele frequency was assessed by measuring the Pearson correlation between the imputed minor allele dose and the true minor allele dose for SNPs binned by minor allele count (Supp. Fig. 2). Comparing dosage in terms of allele count, here, allows for direct comparison across populations that have varying allele frequencies based on the number of individuals. As observed with imputation in other species[34], SNPs with a low number of observed minor alleles in the population have a lower overall imputation accuracy, though imputation accuracy quickly improves with a higher allele count. With an alternate allele count of 8 or more (∼2% minor allele frequency in this population), imputation accuracy was above 90% in most tagging breed groups (Supp. Fig. 2).

## SNP properties of the MNEc2M and MNEc670k arrays

### SNPs in gene coding regions

SNP positions for both arrays were compared to 26,991 predicted and annotated gene models from EquCab2. Of the 2,001,826 SNPs on the MNEc2M array, 591,521(29.5%) SNPs were within 17,128(63.5%) gene models. Likewise, of the 670,805 SNPs on the MNEc670K array, 192,681(28.7%) SNPs were within 14,758(54.7%) gene boundaries. In comparison, of the legacy 74,056 SNPs, 20,950(28%) were within 8,249(30%) annotated gene models.

### SNP informativeness and Inter-SNP distance

Inter-SNP distance was calculated for the MNEc2M, as well as the MNEc670k, SNP sets at different levels of MAF (see Fig. 3 and Table 4) to assess the distribution of SNPs across the genome. On average, 1,250 and 3,756 bp, respectively, separated variants on the two arrays. Informativeness, defined as the number of SNPs with at least one heterozygote, was calculated for the same MAF cut-offs (Table 4). Inter-SNP distance, as well as informativeness, were broken down by breed for both the MNEc2M SNP set (See Supp. Fig. 3) and MNEc670k SNP set (See Supp. Fig. 4).

**Fig. 3.**
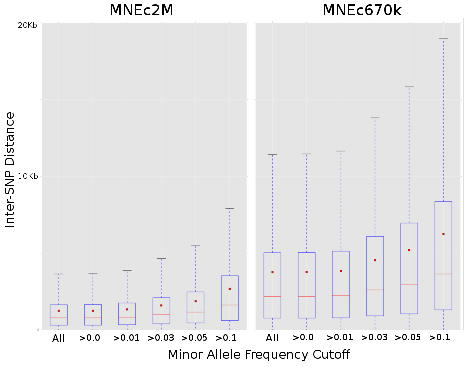
*MNEc2M and MNEc670K Inter-SNP Distance*. Distance between SNPs on each array was calculated using various minor allele frequency (MAF) cutoffs. Considering all available SNPs genotyped on the MNEc2M and MNEc670k arrays, on average, 1,250 and 3,756 bp separate markers (See Table 4 for average inter-SNP distances at the various MAF). Median values (red lines) and mean values (red boxes) were calculated at each MAF cutoff.

### Alternate Allele Frequency

Frequency of alternate (non-reference) SNP alleles were calculated on both arrays using the full 485 sample dataset (WGS+SNP), genotypes derived from whole genome sequence only (WGS Only), and from samples genotyped on the 2M test array described above (SNP Only; see Table S5). Kernel density estimations (KDE) of the alternate allele frequency of SNPs on the MNEc2M array showed a mean frequency between 0.20 and 0.28 (See Fig. 4 and Table 5). In general, regardless of genotyping source, alternative allele frequencies shared similar distributions. Frequency distributions exhibit long tails indicating substantial numbers of samples with non-reference alleles. Alternate allele (ALT) distributions also exhibit a lower median frequency (red line) than mean frequency (red bar), a common property of right skewed distributions [35]. Alternate allele frequency distribution was also broken down by breed for both the MNEC2M array (Fig. 5) and the MNEc670k array (Supp. Fig 5) using genotypes derived from the WGS+SNP sample set. Allele frequency is balanced across breeds, though there were minute differences. For example, the median MAF for Thoroughbreds was 3-12% lower than all other breeds, though this is not unexpected given the reference genome is a Thoroughbred. Despite minor differences, all breeds had long tails and similar distributions of allele frequencies indicating a balanced and representative SNP selection for GWAS.

**Fig. 4.**
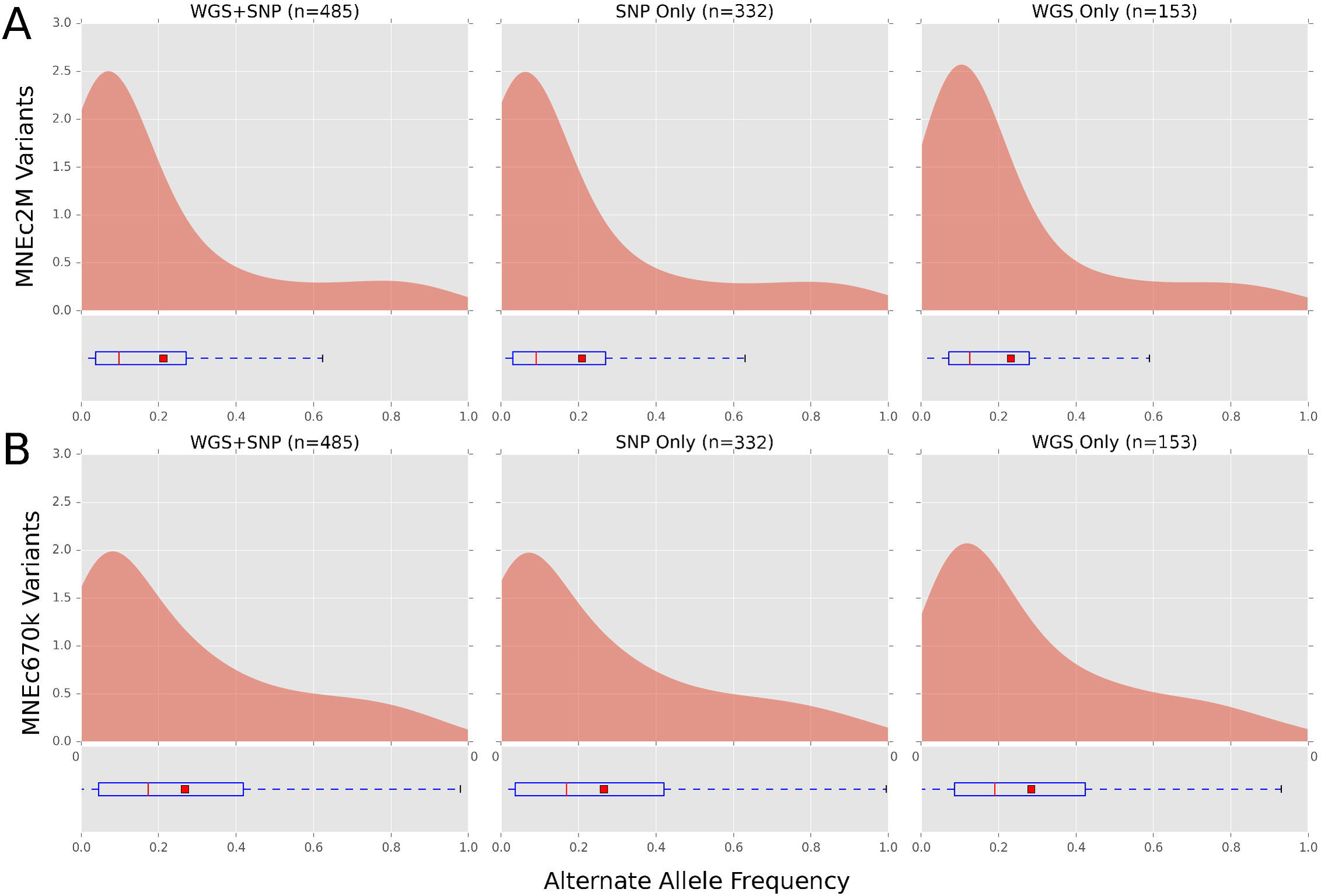
*MNEc2M and MNEc670k Alternate Allele Frequency*. Genotypes from horses on the 2M test array (SNP Only; n=332) as well as whole genome sequence (WGS Only; n=153) were combined (WGS+SNP; n=485) to estimate alternate allele frequency of the SNPs represented on the (**A**) 2M and (**B**) 670K arrays. Fig. 4 shows kernel density estimated (KDE) distributions of alternate allele (ALT) frequency in each sample group using variants that are on each array. Boxplots show ALT frequency distribution median (red line), mean (red square) as well as variance (See Table 5 for values).

**Fig. 5.**
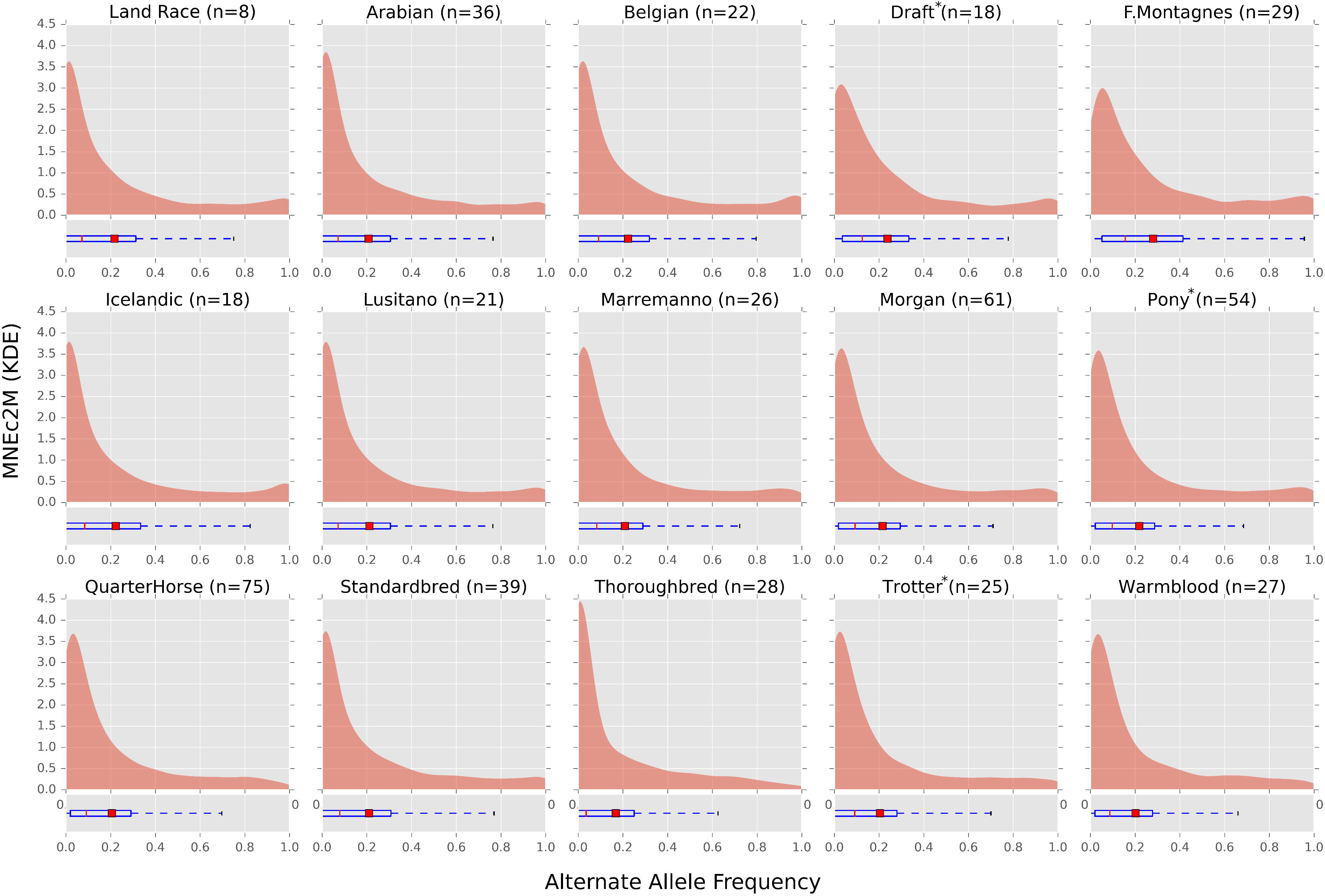
*MNEc2M Breed Specific Alternate Allele Frequency*. Alternate allele frequencies from variants present on the MNEc2M chip were split by breed group. Samples (WGS+SNP) were split into 15 tagging breed groups (See Table S5). Breed groups with asterisk (*) indicate a combination of studbook breeds. Boxplots show median (red line) and mean (red box) of the alternate allele frequency distribution.

### Linkage disequilibrium decay by breed

Linkage disequilibrium was measured using genotype r^2^ between all 1.8M SNPs within 1Mb of one another within and across breeds. LD across breeds (i.e., the WGS+SNP sample sets) (see Fig 6, SNP and WGS curves) decayed faster than LD within any given breed (remaining curves). Within-breed calculations demonstrated that Quarter Horses and the Pony breeds had the lowest LD between SNPs at long distances, decaying to below 0.10 at 1Mb, while Thoroughbreds had the highest LD at all distances considered.

**Fig 6.**
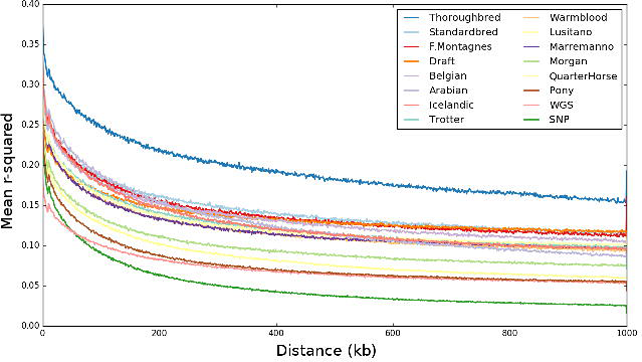
*Linkage Disequilibrium decay within and across breeds*. Pairwise r^2^ was calculated between each SNP within 1Mb having a minor allele frequency greater than 0.05. LD curves are broken down by breed as well as for samples derived from the all-breed WGS as well as from all-breed SNP cohorts. Breeds are ordered (TBLR) in the legend by their r^2^ values at 400kb.

## Discussion

Our goal was to provide a high-quality, standardized SNP array designed for imputation to overcome limitations in SNP density that under-power many genome mapping projects in the horse. To do this, we utilized whole genome sequencing data from 156 horses representing actively-researched and economically-important breeds that were collaboratively collected by 17 laboratories within the equine genetics community. These data, in turn, enabled a large scale, whole genome sequence mapping pipeline with *de-novo* variant calling resulting in a starting set of over 23 million variants. Variant filtering was based on genome coverage, breed representation, linkage disequilibrium, and feasibility of array probe design. This resulted in the successful design of a fully backwards-compatible, high-density, 2 million SNP genotyping array (MNEc2M). Using a test cohort of 485 horses, we identified “tag” SNPs that reconstructed haplotypes both across diverse breeds and within breeds, then selected a subset of ∼670k SNPs for design of the commercially available MNEc670k array. The use of tagging SNPs in this commercial array ensures its utility as an imputation tool up to a SNP density of at least 2 million.

Several successful GWAS in other agricultural animal species have been reported using this same Affymetrix^®^ Axiom^®^ HD genotyping array technology [36–38]. High-density ∼670k genotyping arrays were used in domestic cattle to identify structural variation [39,40]. Other studies have leveraged this technology to discover and refine loci associated with production traits [41–43]. Based on reports from other domestic animals, coupled with preliminary analyses performed here, we anticipate similar performance and increased power for map-based studies to soon follow in the horse.

### Assembly correction of the equine genome

Initial filtering for array compatible SNPs immediately reduced the pool from ∼23 million SNPs that were discovered from *de novo* variant calling to 10 million SNPs compatible with array design. These purely technical constraints, such as removing all variants that were in close proximity to one another, substantially reduced the amount of variants considered here. While we thoroughly characterized the SNPs that were ultimately included on the genotyping array, the entire 23M SNP set has been submitted to dbSNP and the European Variation Archive. Additional analyses are being performed to further enhance knowledge of equine genetic diversity at the WGS level. This includes investigation of the > 400,000 sites where all WGS alleles differed from the reference allele determined by Sanger sequencing of Twilight. If the discrepancies in these data are verified, this information will be useful for further annotation and error correcting of the current equine genome reference assembly as well as for future genome assemblies. Here, we focused on variants that were compatible with array design, however, we anticipate that improved genome annotation and finer scale haplotype maps will result from this larger SNP set.

### Determining highly precise, gold standard SNP sets

Quality control and evaluation of conversion rates on the previous arrays have shown that many of the SNPs assayed on the legacy arrays are non-informative or not polymorphic in many breeds (See Table S2). These SNPs could truly occur at low frequencies within those populations, could have been singleton mutations in the individuals included in the legacy SNP discovery panel, or may have been false positive sites where no true genomic variation occurs. Compared to the SNP discovery panel used in the legacy arrays, the cohort of 153 horses used here provided a valuable opportunity to identify a precise set of SNPs that would be informative across many breeds.

While variant discovery from WGS had substantially higher overall depth of coverage than the SNP50 discovery effort, many breeds still had a modest number of individuals or relatively low average depth of coverage (Table S1). Still, genotype comparisons at variant sites from WGS to legacy arrays showed an overall high congruency rate (94% at 6X and 99% at 12X; Fig. 1), however, many SNPs with high QUAL score did not agree between WGS and legacy arrays. Since we were unable to determine the source of error in genotype mismatches between the WGS discovered variants and the legacy SNPs, especially those with high QUAL scores, we used additional information such as how many variant calling groups a SNP was discovered in as well as sequencing read level information to determine the criteria for including WGS discovered SNPs on the MNEc2M and MNEc670k arrays.

Distinguishing marginally higher quality variants, even at overall validation rates of over 94% becomes important when selecting a small fraction of variants to be included in a commercial array. Selecting a final subset (670k) containing only 6% of the initial ∼10 million array-compatible variants posed an opportunity to control for overall false discovery rate during SNP selection. Variant recalibration allowed successful identification of variants with high QUAL scores that were less discordant between WGS and SNP50 genotypes. In the absence of high-confidence training SNP sets (i.e., dense HapMap data), we defined several different “gold standard” SNP sets for variant recalibration. While variant recalibration performed better than QUAL scores alone, there was not a single training set that clearly out-performed the others (Fig. 2). However, using recalibrated VQSLOD scores coupled with a focus on identifying high precision SNPs, did result in a high conversion rate of probes (92.3%) on the 2M test array. As more WGS data are generated in more horses allowing for further comparisons between SNPs genotyped using different technologies, and SNPs discovered here can be further validated, the ideal SNP training set for variant recalibration will become more formalized.

### Genotyping success in blood versus hair root DNA

DNA samples genotyped on the MNEc2M array were primarily derived from blood, but also came from hair roots. Both DNA sources had failed samples, however, a substantially higher fraction of hair root samples produced poor genotyping rates. Hair root DNA also tended to have lower DNA concentrations when samples were re-hydrated. To maximize information, we chose to genotype all submitted samples, even though samples from hair roots did not meet minimal DNA quantity guidelines specified by Affymetrix, though samples were dropped if they did not meet genotyping quality control thresholds.

To determine if sample origin (blood versus hair roots), DNA quantity, or both, were associated with failure to pass genotyping quality control metrics, a logistic regression for sample success on DNA concentration (Pico Green) and blood/hair status was performed (Supp. Fig. 6). It is clear that higher DNA concentrations increased the probability of genotyping success. However, while blood samples were more likely to produce passing samples regardless of DNA concentration, at adequate concentrations, hair root samples were also highly likely to produce passing genotypes.

We also noted variation in genotyping quality of specific SNPs based on tissue origin. During array design, SNPs were not disqualified that potentially performed better in blood versus hair root, e.g. ‘PolyHighResolution’ in one and not the other (Table S4). While it is difficult to determine if certain probes perform better using DNA isolated from one tissue versus the other, in the samples tested here, DNA isolated from blood was preferred over hair root when DNA concentration is low or questionable.

### Setup for precision imputation

Although the legacy equine arrays were not designed for imputation, several previous studies have demonstrated their utility in imputing genotypes, which, if performed on a larger scale across many breeds, could greatly improve the chances of success in horse genome mapping studies. A study in Standardbreds, Quarter Horses and Thoroughbreds showed high fidelity imputation from legacy arrays (54k and 65k) to a higher marker density (74k) using a reference population of 248 horses [3]. Another study found that genotype imputation in Thoroughbreds was feasible from a very low density (1-3K markers) to a legacy SNP set (70K), although it was impacted by the minor allele frequency of the SNPs being imputed, as well as potentially complex LD structure [44].

We masked variants genotyped on the MNEc2M array down to the MNEc670k SNP set in a test cohort of 485 horses and found a high overall imputation accuracy (96-99%) up to the 2M density, across several different breeds of horses (See Table 3). It is important to note that the 96% average imputation accuracy in the Draft horses was mainly due to underperformance (92.6%) in the Percheron horses. This is not a surprising result as Percherons were underrepresented in the Draft breed group (2 out of 18 individuals) thus removing both horses from the imputation reference population for validation yielded poor results. The inclusion or exclusion of samples in the reference population can significantly impact imputation accuracy. Here, it was critical to maximize the available data during the SNP discovery process, especially for those breed groups that had limited sample representation.

While our imputation scenarios focused on imputing to the MNEc2M SNP set, imputation to higher SNP densities (>2M) is feasible. A recent study using whole genome sequencing from 44 Franches-Montagnes and Warmbloods imputed SNPs from the legacy SNP50 genotypes to nearly 13M variants with 95% accuracy [45]. WGS breed representation in this study varied between 1 and 29 individuals. In the future, it will be critical to expand the number of WGS samples in the reference population described here in order to attain proper breed representation for reliable imputation to higher SNP densities.

As sample size continues to plague the characterization of complex equine phenotypes, despite falling genotyping costs, the development of this affordable, backwards compatible, 670K imputation array will allow the inclusion of already genotyped individuals in higher-powered analysis, providing an economical trade-off for studies that must choose between more markers or more samples. We expect that imputation will complement future GWAS studies and mapping studies designed based on haplotype structure. In the future, improvements in imputation software as well as development of equine-specific imputation protocols that leverage breed information (e.g. recombination maps) will further increase accuracy of genotype imputation. Furthermore, as additional WGS are integrated into reference populations, imputation performance will continue to improve.

## Conclusions

Here we report, through a community effort, the leveraging of WGS data from 153 individuals with a 13X median coverage to achieve new variant discovery in the domestic horse. The use of WGS enabled us to generate orders of magnitude more data than previous technologies allowed. Empirical SNP properties were used to produce composite scores for each SNP based on machine learning approaches, allowing better detection of false positive variant calls that could not be achieved using generic QUAL scores alone. Thus, from a starting set of over 23 million SNPs, we identified a set of ∼5 million SNPs which were considered for array design. With probe design recommendations from Affymetrix, we further filtered this list to 2M SNPs (MNEc2M), which adequately represented the breeds used in variant discovery, were evenly spaced across the genome, and had the highest chance of conversion in the array implementation.

Using a test cohort of 332 horses, we used the MNEc2M SNP set to identify haplotype tagging SNPs both within and across breeds. Choices made in 2M array design, particularly with regard to variant filtering, resulted in greater than 92% of the SNPs on the 2M array (MNEc2M) returning high-quality genotypes. Filtering further, we designed the MNEc670k genotyping array to contain SNPs which allowed for accurate imputation. Together these genotyping platforms represent the next generation of genomic array technologies for the domestic horse.

## Methods

### Whole-genome sequencing and mapping pipeline

Paired-end Illumina Hi-Seq 100 base pair whole genome sequences were generated for 153 horses representing 24 distinct breeds (Table S1). Raw reads for each individual were mapped using the PALEOMIX pipeline [26] and aligned to an extended EquCab version 2.0 genome (see below). Specifically, reads for each individual were processed separately by sequencing lane and filtered for quality control using the program Adapter Removal [46], which removed PCR adapters, trimmed low quality base-pairs, and collapsed overlapping paired-end reads into a single high-quality read, and the resulting reads were filtered by length. Passing reads were mapped to each reference file using BWA [47]. Paired-end reads for which the mate was filtered were mapped in single-end mode. PCR duplicates were detected and removed, the resulting bam-files were merged, and reads were realigned around detected indels.

### Extended EquCab 2.0 reference genome

An extended version of the EquCab2 reference genome was used which included the 31 autosomal chromosomes, ECAX, and equine chromosome unknown 1 (ChrUn1) [1], together with an additional 7,850 contigs designated as ChrUn2 generated from unmapped Twilight genomic DNA reads which were de novo assembled using the Velvet assembler [48]. These additional contigs were required to meet any of the following criteria: 1) longer than 1,000 base pairs with no BLAST alignment (bit score > 99) to the human, canine, or bovine genomes; 2) longer than 1,000 base pair and either a single BLAST alignment to a human, canine, or bovine chromosome or multiple alignments that mapped to a single chromosome; 3) between 500 and 999 base pairs and a BLAST alignment to a single human, canine, or bovine chromosome with a bit score >499; 4) between 500 and 999 bp with alignment to a single chromosome where the total coverage of the aligned region included more than 80% of the coding length of an existing human, canine, or bovine annotated protein-coding gene.

### PALEOMIX *vcf pipeline* implementation and python source code

Programs within PALEOMIX are abstracted as nodes within the program to allow files to be generated within a temporary directory and only to be moved to the final directory upon successful completion. In addition to nodes for the read alignment mapping programs, additional nodes were created to run variant calling programs implemented by GATK [27] and SAMtools [28]. Additional PALEOMIX nodes were created to perform variant recalibration as well as to assess precision versus recall for Morgan and Standardbred breed groups (Fig. 1 and Fig. 2). Source code for the extended PALEOMIX nodes are available at https://github.com/schae234/pypeline.

### Variant calling and validation PALEOMIX nodes

The PALEOMIX computational framework was further extended to process alignments and produce variant calls. Individuals were split into 16 different calling groups based on both breed and sequencing depth in order to minimize biases due to coverage and population stratification. Variant call files (.vcf) were produced for each group using both GATK Unified Genotyper with” –stand_call_conf” set to 30, “—stand_emit_conf” set to 10, and “–dcov” set to 200. Variants called by GATK were retained if they were independently called by SAMtools “mpileup” with default calling parameters. This allowed possible false negatives to be assessed downstream using other quality metrics. Of the 26,884885 SNPs called by SAMtools and 31,506,364 SNPs called by GATK, 22,557,988 variants were called independently by both callers and retained for further analysis.

### Variant recalibration (VQSLOD scores)

Both GATK UnifiedGenotyper as well as SAMtools assign generic quality scores (QUAL) to each discovered variant, which is the posterior probability that a true variant exists given the pileup of reads at a given locus using base pair quality and expected allelic distribution of samples. Using GATK, additional metrics were generated for each variant including coverage (DP), quality of depth (QD), Fisher strand bias (FS), mapping quality rank sum (MQRankSum) and read position rank sum (ReadPosRankSum). Definitions from the GATK manual for each metric are below.

**Coverage (DP)** – Total, unfiltered depth of coverage.

**Quality by Depth (QD)** – Variant confidence (from QUAL field) divided by depth of non-reference samples.

**Fisher Strand Bias (FS)** – Measure of strand bias, i.e. the variation seen on only the forward or reverse strand.

**Mapping Quality Rank Sum (MQRankSum)** – The rank sum test for mapping qualities.

**Read Position Rank Sum (ReadPosRankSum)** – The rank sum test for the distance of the variant from the end of reads.

These parameters were used to train linear mixed models using GATK VariantRecalibrator which produces a composite variant quality metric called a VQSLOD score which can be summarized as using the formula VQSLOD ∼ DP + QD + FS + MQRankSum + ReadPosRankSum. We defined three “gold standard” datasets: 1) SNPs previously genotyped on the 54K SNP chip, 2) variants called in four or more calling groups, 3) and variants within the 99^th^ percentile of QUAL scores. Models were trained on chromosomes 2-31 and validated on chromosome 1 in both Morgan and Standardbred breed groups as a subset of each breed group had whole genome sequence data at target depths of 6X (12 Morgans and 6 Standardbreds) and 12X (6 Morgans and 4 Standardbreds) target coverage (actual coverage MOR6X: 5.23-6.96X, mean=6.11X; STD6X: 4.61-5.89X, mean=5.29X; MOR12X: 10.42-15.14X, mean=12.96X; STD12X: 11.21-11.99X, mean=11.63X). These “gold standard” training SNPs were used to assess the precision of different quality metrics of *de novo* variant calling.

### Preliminary 5M high-density SNP selection

A preliminary list of approximately 5 million SNPs was prepared for probe design (conversion) recommendation by Affymetrix. VQSLOD scores were generated for all ∼23 million initial bi-allelic candidate SNPs. SNPs were filtered out if they 1) fell within repetitive regions designated by RepeatMasker 3.3.0 [49] (780 SNPs); 2) fell within 20bp of another variant (9,342,733 SNPs); and 3) had fewer than 2 observed instances of minor allele (4,858,273). Of the remaining 11,435,936 SNPS, further filtering was applied to obtain equal coverage throughout the genome. Pairwise LD was calculated between SNPs within 10kb windows. If SNPs within a window had an r^2^ value of over 0.90, priority was given to SNPs that were in called in Draft or Pony groups. If a SNP was in an equal number of priority groups, VQSLOD scores were used to break ties. After all filtering criteria were applied 5,443,950 SNPs remained.

### High-density 2M SNP selection

From the ∼5 million candidate SNPs, Affymetrix provided four classes of recommendations based on the probability the SNP would be convertible through probe design (‘recommended’, neutral, ‘non-recommended’, ‘not possible’). A recommended SNP has: a probability of conversion > 0.60, no interfering polymorphisms within 24 bases, and unique flanking sequence. Non-recommended SNPs have: either duplicate flanking sequence, a probability of conversion < 0.40, interfering polymorphisms within 21 bases, or more than 2 interfering polymorphisms within 24 bases. A ‘not possible’ designation is given to probes which probes cannot be created, and neutral recommendations cover all other cases. Furthermore, SNPs with alleles of A/T or C/G required two probes to differentiate between allele states and were tagged as allele-specific SNPs. Recommendations were generated for both forward and reverse strands based on the above criteria. Groups of SNPs were ranked within 50kb windows and probe design criteria were chosen using a greedy algorithm until 37 SNPs were chosen per window using the following criteria: 1) VIP SNPs previously designed on 54/65K chip (regardless of recommendation or strand), 2) SNPs for known Mendelian traits such as coat color (regardless of recommendation or strand), 3) SNPs designated as ‘recommended’ by Affymetrix and designable with one probe, 4) SNPs with a ‘neutral’ recommendation from Affymetrix designable with one probe, 5) any SNP within the equine MHC region, 6) SNPs requiring multiple probes. If a SNP was equally designable in forward and reverse strand, the forward strand was chosen for interpretability. Using these criteria, 2,001,826 SNPs were chosen for array design.

### 2M SNP test array and sample quality control

Using probes designed according to the above criteria, 347 horses from 20 breeds were genotyped on the 2M array using DNA isolated from blood (n=286) using the Gentra PureGene Blood kit. DNA from hair roots (n=61) was isolated using the Gentra Puregene Blood Kit with a modified version of the Gentra Puregene Mouse Tail purification protocol. The amount of Proteinase K was increased to 20 µL, isopropanol to 650 µL, and ethanol to 500 µL. The DNA hydration solution was reduced to 20-30 µL depending on size of DNA pellet. Quality control metrics were calculated on arrays grouped by tissue type (blood vs hair) as well as array batches, and arrays were dropped for various reasons at multiple QC steps. “DishQC” evaluates call rates between A/T probes and C/G probes of non-polymorphic sites to assess background probe contrast and was calculated using Affymetrix Power Tools. Samples with DQC scores below 0.60 were removed from further evaluation. Passing arrays were then genotyped using approximately 20,000 high-confidence probes provided by Affymetrix in their R1 release package (See https://www.thermofisher.com/order/catalog/product/550583#/legacy=affymetrix.com). If the number of arrays per tissue/array group was above 96 samples, generic priors were used for probe-set genotyping; otherwise, SNP specific priors were used (provided in the R1 package released by Affymetrix). Sample arrays with 20K call rates below 0.97 were removed from further analysis.

Remaining samples (n=332) were genotyped on all probes using a similar approach. Generic or SNP specific priors (provided in R1 package) were used for groups containing more or less than 96 samples, respectively, and samples with call rates below 0.97 were removed from the analysis. Successfully genotypes samples were then grouped by genotyping plates to check for batch effects. Plates with pass rates below 95% in blood or 93% in hair were dropped.

Up to 4 probes were used to genotype each SNP across samples passing quality control criteria. Probe performance was calculated using the `*Metrics.R`* script provided by Affymetrix which assessed several quality criteria. Twelve criteria were used to assign each SNP into 6 categories representing probe conversion types. High-quality probes fell in one of three categories in descending order of quality: ‘PolyHighResolution’ SNPs had good cluster resolution and at least two examples of the minor allele, ‘MonoHighResolution’ SNPs had good SNP clustering but less than two samples with the minor allele, and ‘NoMinorHom’ had good cluster resolution but no samples with homozygous, minor alleles. Poor quality SNPs were qualified as having off target variation (OTV), a call rate below threshold, or a combination of poor performing cluster properties. SNPs with multiple probes were assigned a best probe based on the above high-quality conversion types with the added constraint that SNPs with no minor homozygote did not result in extreme Hardy-Weinberg values (p ≤ 10e-5; Chi-Squared Test). 1,846,988 SNPs passed the above quality control metrics.

### 670K commercial array SNP selection

Genotype information derived from the horses on the 2M test array (n=332) were combined with SNPs in horses called by whole genome sequencing (n=153) resulting in a total of 485 horses genotyped at 1,846,988 SNPs. Tag SNPs were calculated using FastTagger [32] using two different tagging scenarios. Tagging SNPs informative across populations (inter-population) were identified by running FastTagger on the entire dataset. Parameters provided to FastTagger identified inter-population tag-SNPs at a resolution of down to 0.01 minor allele frequency and up to 0.99 r^2^.

To identify SNPs tagging population specific haplotypes (intra-population), samples were split into 15 tagging breed groups based on available sample size: Land Race, Arabian, Belgian, Draft, Franches-Montagne, Icelandic, Lusitano, Maremanno, Morgan, Pony, Quarter Horse, Standardbred, Thoroughbred, Trotter, and Warmblood (See Table S5). FastTagger was run with parameters to differentiate tag-SNPs within each population at a resolution of 0.10 minor allele frequency and 0.90 r^2^. Representative SNPs from each population specific set of tag SNPs were collapsed using the program `Multi-Pop-TagSelect` which assesses overlap between sets of tag SNPs and identifies the subset of SNPs which near-minimally spans the set of populations [33]. Intra-breed specific SNPs were kept as long as they were tagged haplotypes in five or more breed groups.

Quarter Horses, Ponies, Morgans, and Standardbreds needed a much higher number of intra-breed specific SNPs to tag haplotypes, indicating they were the most diverse breeds. Additional tag SNPs (n=13,993) were included in the final commercial array if they tagged haplotypes in three or more of these diverse breed groups.

### Genotype imputation between the MNEc670k and MNEc2M arrays

A reference population of the 485 horses with genotypes at ∼2M SNPs (generated either by the MNEc2M array or whole-genome sequence) was used as a reference population for imputation. Horses from each different breed were used to test the imputation accuracy of the array. A random set of 1/3 of individuals were removed from each breed group and masked down to the MNEc670k SNP set as well as the SNP65 SNP set. Genotypes were then imputed using the remaining (non-masked) individuals as a reference population. Imputation was performed using Beagle 4.0 [50] using default parameters. Genotype concordance was calculated using VCFtools (0.1.15) using the `--diff-indv-discordance` option [51]. Briefly, concordance, as calculated by VCFtools, is the proportion of non-missing SNPs in both datasets that have the same allele calls. Missing genotypes are not considered in the concordance calculation and phase is not taken into account (e.g. A/T and T/A are concordant). VCFtools reports the proportion of matching genotypes per individual to the number of sites that are shared across input SNP sets.

## Abbreviations

SNP: Single Nucleotide Polymorphism
MNEc2M: the 2 million SNP array developed here
MNEc670k: the 670 thousand SNP array developed here
WGS: whole genome sequencing
bp: base pair
GATK: genome analysis tool kit
QUAL: generic quality score output by GATK and SAMtools
MOR: the Morgan horse breed
STD: the Standardbred horse breed
VQSLOD: variant quality score log-odds
Gb: giga-bases
MHC: major histocompatibility complex
ECA: Equus caballus chromosome
VIP: very important probe
MAF: minor allele frequency
ALT: alternate allele
KDE: kernel density estimation
LD: linkage disequilibrium
ChrUn1: Equus calballus unknown chromosome

## Declarations

### Acknowledgements

These genomic tools are the result of a collaborative effort that involved a number of laboratories within the equine genetics community, many of which provided support or insight at various steps for this community resource. We would like to acknowledge Chad Dow from Affymetrix for the support he and his team provided while designing the MNEc2M array.

### Author Contributions

Experimental concept and design: MEM, JRM, LO, RJS, MS; Sample collection and data contribution: EB, DLB, EB, GKB, GB, SAB, OD, RF, CJF, VG, BH, VJ, TK, TL, GL, MSL, NM, ACM, JNM, AM, JM, CP, SP, SR, IT, JT, GT, AVS, CMW, BW, LO, JRM, MEM; Data analysis and interpretation: RJS, MS, MEM; Computational support: RJS, MS; Manuscript writing and figures: RJS, JRM, MEM; Manuscript review: All authors read and approved the final manuscript.

### Animal Welfare

DNA samples were previously collected with approval from the Animal Care and Use Committees at the respective institutions.

### Data Availability

Whole genome sequences are available in the following NCBI BioProjects: PRJEB14779, PRJNA273402, and PRJEB10098. Additional sequences have are restricted in availability due to preexisting material transfer agreements and can be requested by contacting the contributing investigator in Table S1. Genotypes for horses on the MNec2M array will be released upon publication. Genome positions for all 23 million discovered SNPs have been submitted to dbSNP as well as the European Variation Archive.

## Tables

**Table 1**

**SNP Sets used at various steps in array design**

Variants discovered from whole genome sequencing were filtered at various steps for quality control or using array design criteria. Six distinct sets of variants ranging from the initial ∼23M high-quality, variants discovered from WGS to the 670k variants available in the commercial genotyping array are described throughout this manuscript.

**Table 2**

**Number of breed specific tagging SNPs**

Number of tagging SNPs required to reconstruct haplotypes in each breed (MAF > 0.10 and 0.90 r^2^, See Methods).

**Table 3**

**Imputation accuracy of the MNEc670k SNP genotyping array**

Breed specific imputation accuracy (mean +/- s.e.m.) of genotypes from MNEc670k to MNEc2M SNP sets. In each tagging breed group, 1/3 of samples genotypes were masked to lower density SNP sets and removed from the reference population of 485 horses. Imputation was performed using Beagle 4.0 and concordance was determined with VCFtools.

**Table 4**

**MNEc2M and MNEc670k Inter-SNP distance at various minor allele frequency cutoffs**

Inter-SNP distance was calculated between SNPs informative at minor allele cutoffs greater than 0, 0.01, 0.03, 0.05 and 0.10. The number of SNPs included at this MAF cutoff is included. Distance and informativeness was re-calculated on both MNEc2M and MNEc670k arrays which were further broken down by tagging breed group (See Table S5).

**Table 5**

**MNEc2M and MNEc670k variant mean and median alternate allele frequency**

Average (mean and median) values for MNEc2M and MNEc670k arrays broken down by genotype information available from WGS, CHIP or WGS+CHIP.

## Supplementary Files

**Table S1**

**Whole genome sequencing (WGS) samples**

Whole genome sequencing samples with horse identification and read depth. 153 Individuals representing 24 breeds (Twilight is included as 4 entries). Table includes horse identifier, breed, contributing laboratory and coverage statistics for each individual in both the nuclear and mitochondrial genomes. Variant calling groups indicate which individuals were grouped together during variant discovery.

**Table S2**

**SNP50 breed informativeness**

Impact of breed on SNP50 informativeness. Table shows informativeness (MAF ≥ 0.05) of the legacy, SNP50 SNP set for light, draft, and pony breeds described by McCue et al. [1,2]. Mean number of informative SNPs per breed was ∼42,000 (∼77%). 19,427 SNPs were informative in all 14 breeds.

**Table S3**

**SNP validation by discovery by variant calling group**

SNPs discovered from whole genome sequence (See calling groups in Table S1) were validated by assessing the minor allele frequency in individuals genotyped on the MNEc2M test array.

**Table S4**

**MNEc670k SNP information**

Contains information on MNEc670k SNPs. Includes Affymetrix probe ID, genetic coordinates, MNEc Identifier, tissue origin indicating high quality genotyping in either hair root or blood, Affymetrix assigned genotyping cluster resolution, and inclusion criteria:

Intra – SNP tags intra-breed haplotype

Inter – SNP tags inter-breed haplotype

MHC – SNP within Equine Major Histocompatibility Complex region

Diverse – SNP tags haplotype in 3 or more of diverse breeds that need many tagging SNPs (Quarter Horse, Pony, Morgan, Standardbred; See Table S4)

Density – SNP included to target at least 8 SNPs per 50kb genomic window (targeting uniform genomic coverage)

**Table S5**

**Sample breed and tagging breed groups**

Samples (n=485) from either whole genome sequence (WGS) or from the 2M test array (SNP) were assigned to one of 15 tagging breed groups based on their reported breed.

**Supp. File 1**

**Position of variants discovered from WGS data**

File contains chromosome, bp position, distance from previous discovered SNP and alternate allele frequency for the 23 million SNP set.

**Supp. File 2**

**MNEc2M SNP information**

Contains information for SNPs included on the MNEc2M array. Columns include MNEc identifiers, EquCab 2.0 coordinates, and fields indicating if the SNP was discovered in Draft and Pony groups (which were under-represented on the legacy arrays, See Table S2).

**Supp. File 3**

**Breed specific tagging SNPs**

File containing informative tagging SNPs broken down by breed.

## Supplementary Figures

**Supp. Fig. 1.**
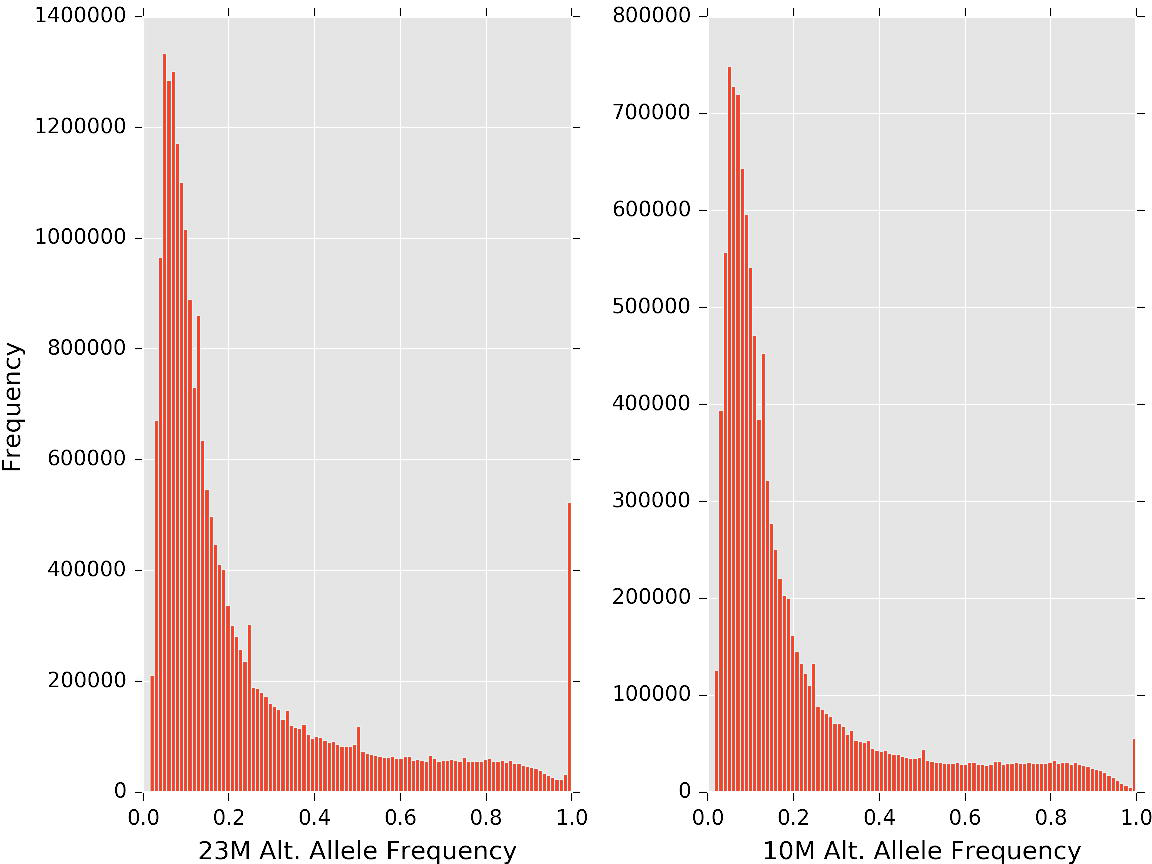
Alternate allele frequency for 23M and 10M SNP sets. Histograms show the alternative allele frequency for the ∼23 million SNPs discovered by WGS and the ∼10 million SNPs that were compatible with array design (see Table 1). A high number of SNPs were observed at 100% alternate allele frequency in a sample cohort that included deep sequencing (see Table S1) of Twilight (the horse used for the reference genome) indicating likely Sanger sequence errors.

**Supp. Fig. 2.**
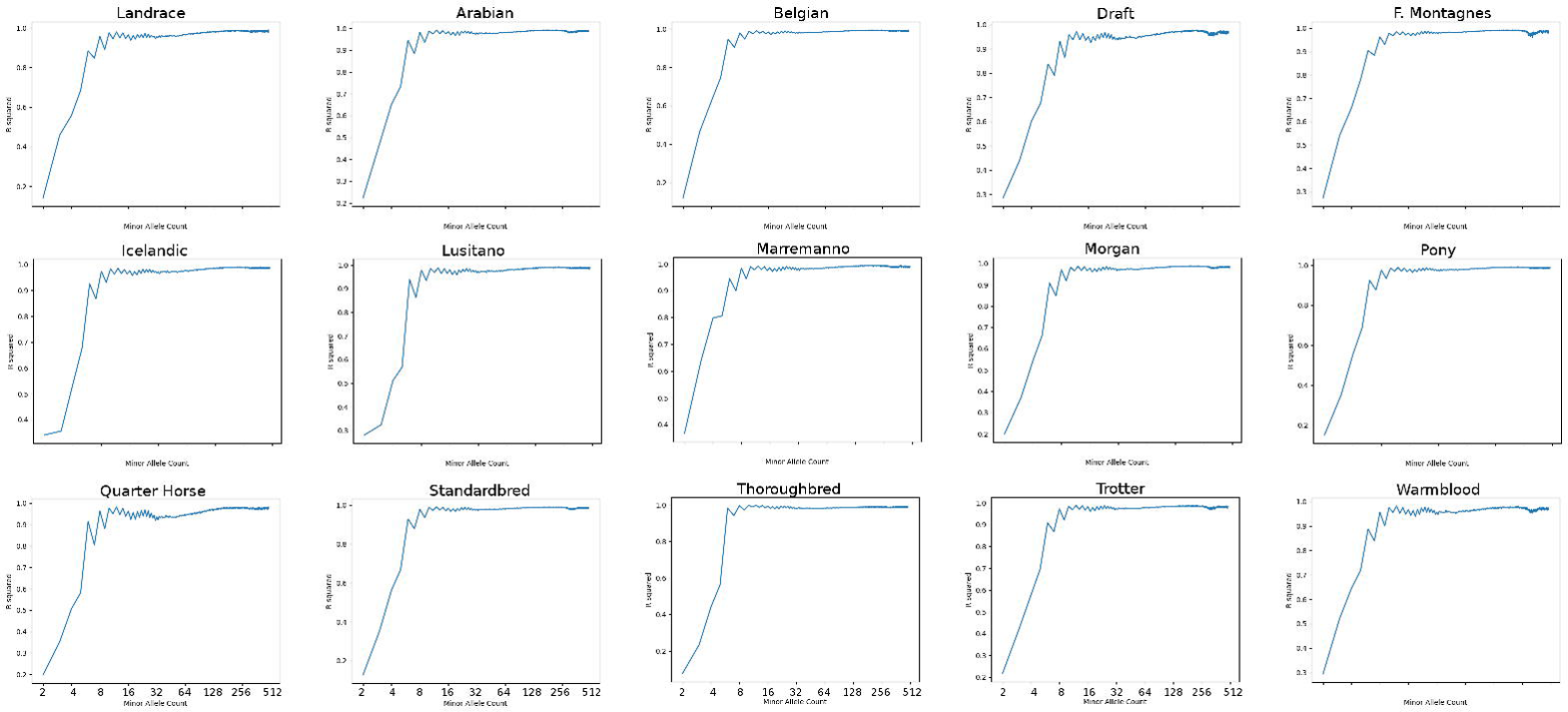
The effects of minor allele on imputation accuracy. SNPs were binned by minor allele count of the marker in the reference panel for each tagging breed group. SNPs were masked down to the 670K set then imputed up to ∼2M. For SNPs in each minor allele count bin, a Pearson correlation was calculated between the imputed minor allele dosage and the true minor allele dosage. The x-axis in each panel are on a log scale to show the relationship for low minor allele count SNPs.

**Supp. Fig. 3.**
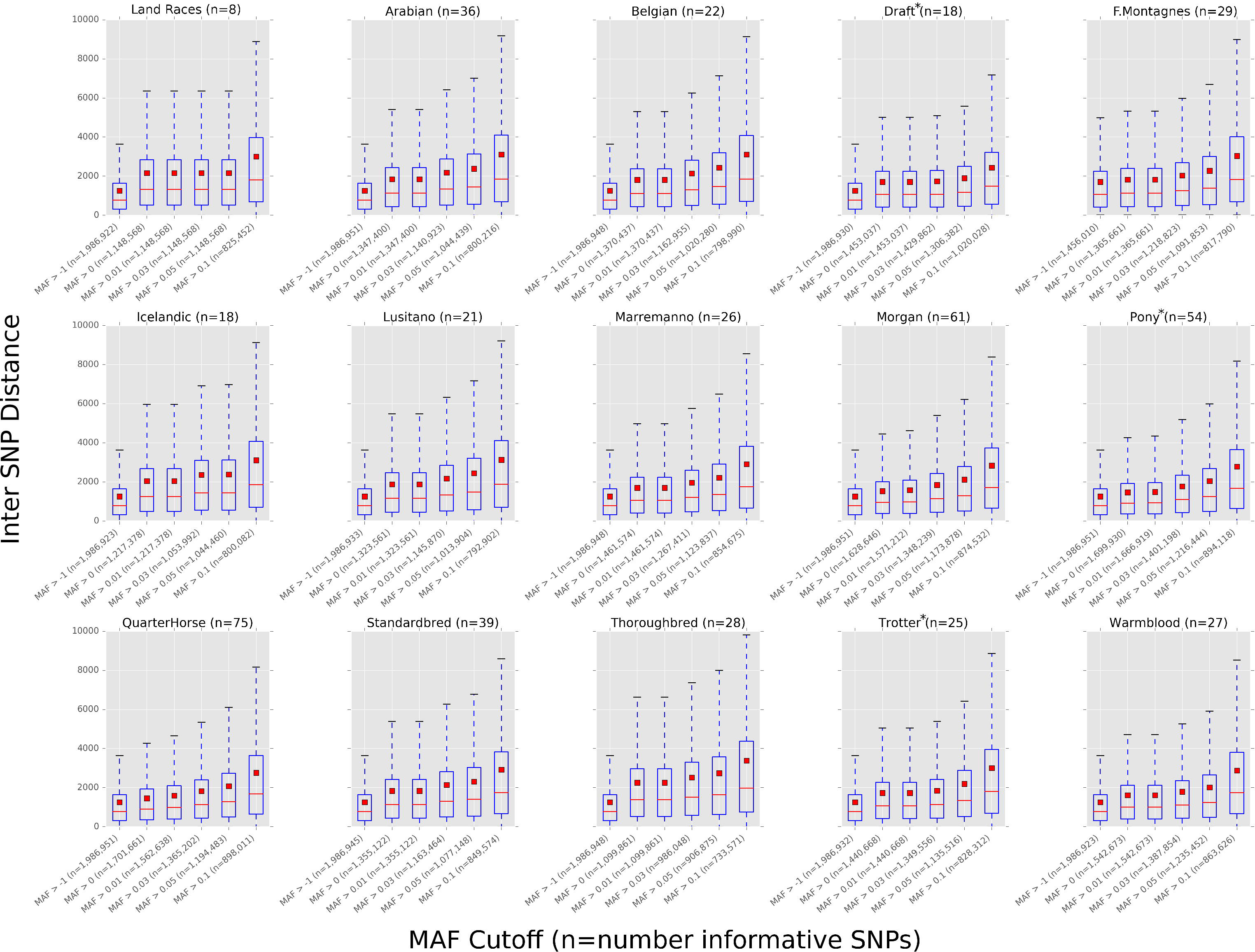
MNEc2M Inter-SNP distance and informativeness by breed. Distance between SNPs on the MNEc2M array were calculated by breed group (See Table S5) at various minor allele frequency cutoffs (0, 0.01, 0.03, 0.05, 0.10). Breed groups with asterisk (*) indicate a combination of studbook breeds.

**Supp. Fig. 4.**
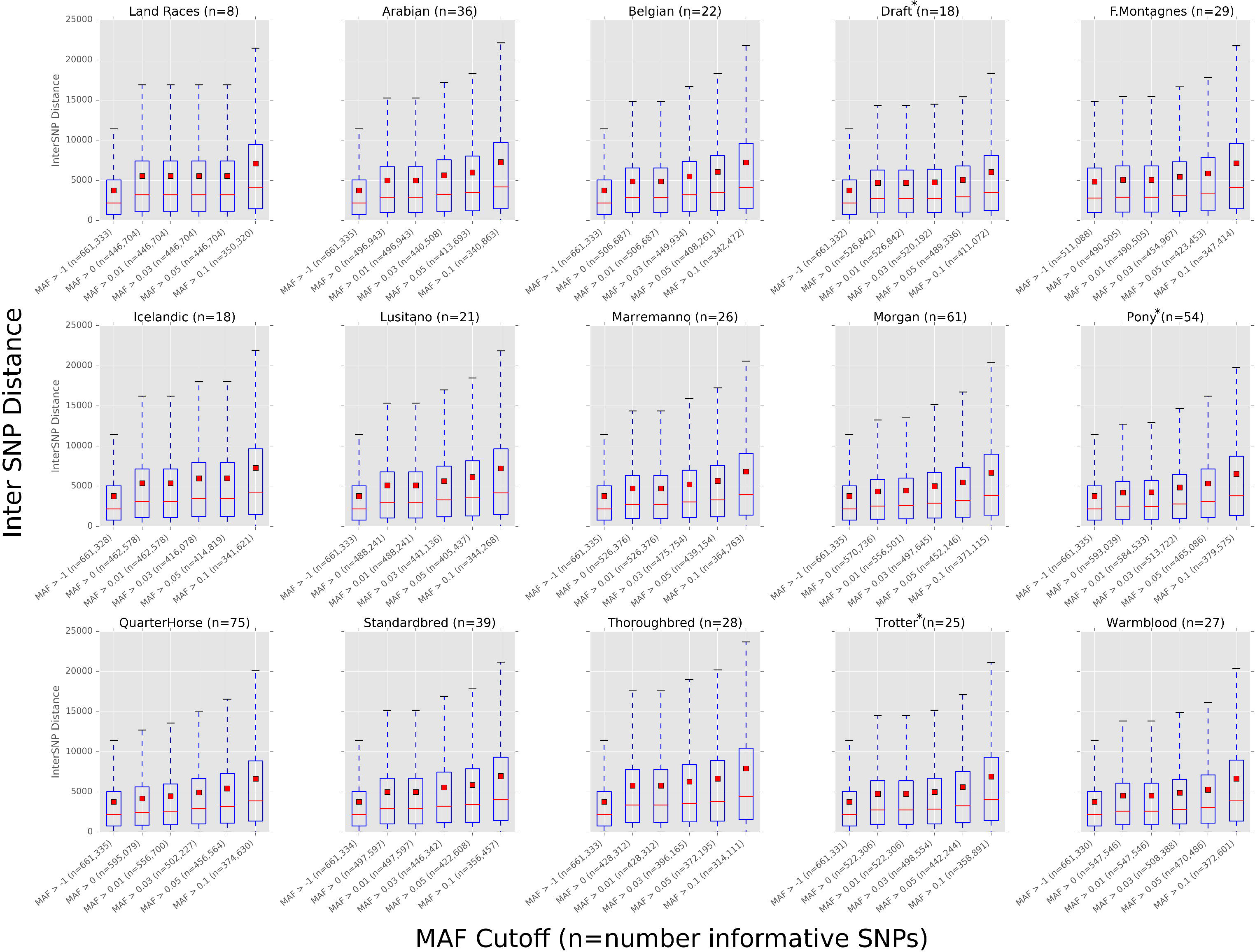
MNEc670k Inter-SNP distance and informativeness by breed. Distance between SNPs on the MNEc670k array were calculated by breed group (Table S5) at various minor allele frequency cutoffs (0, 0.01, 0.03, 0.05, and 0.10). Breed groups with asterisk (*) indicate a combination of studbook breeds.

**Supp. Fig 5.**
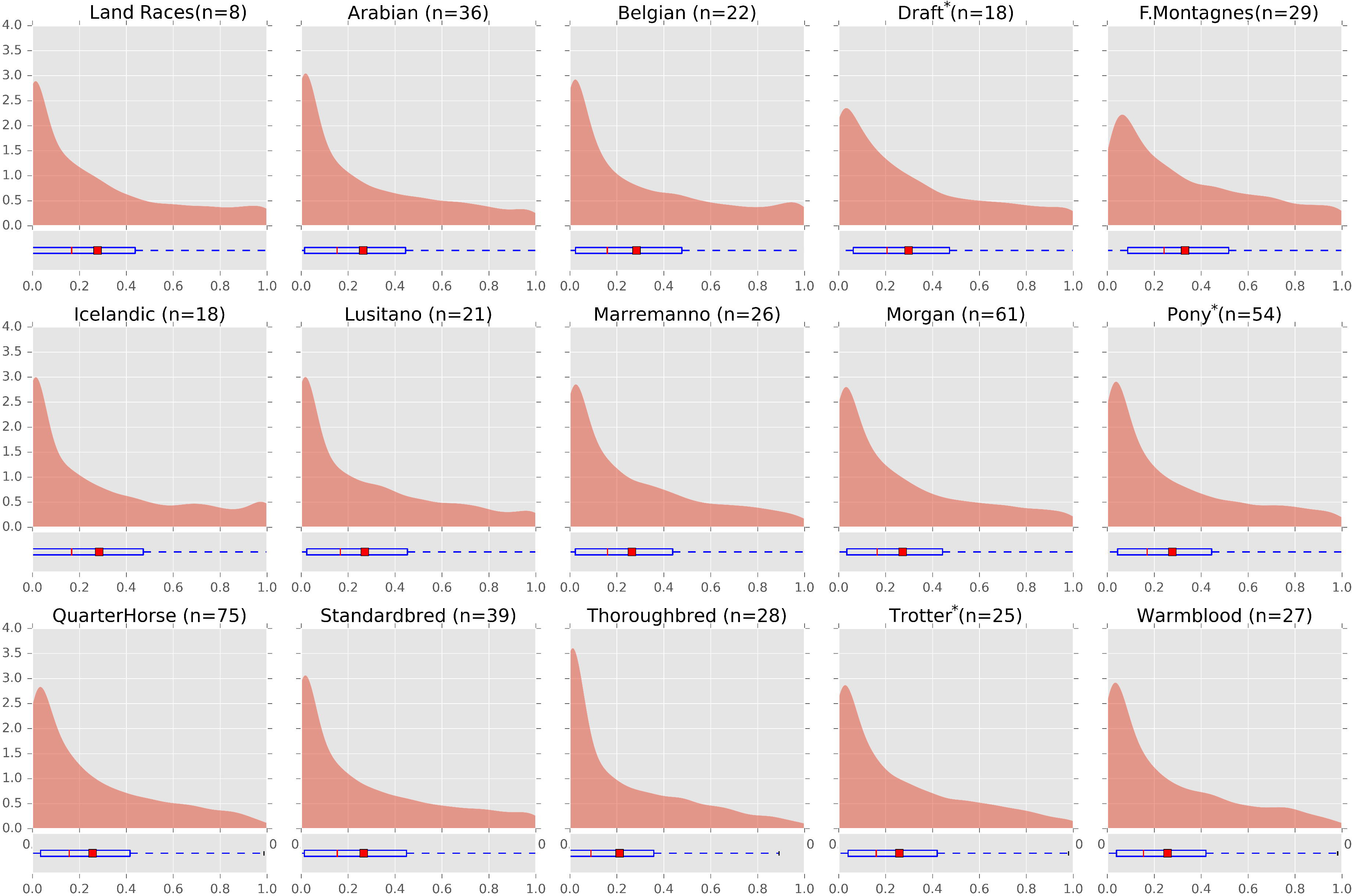
MNEc670k Breed Specific Alternate Allele Frequency. Alternate allele frequencies from variants present on the MNEc670k chip were split by breed group. Samples (WGS+SNP) were split into 15 tagging breed groups (See Table S5). Breed groups with asterisk (*) indicate a combination of studbook breeds.

**Supp. Fig. 6.**
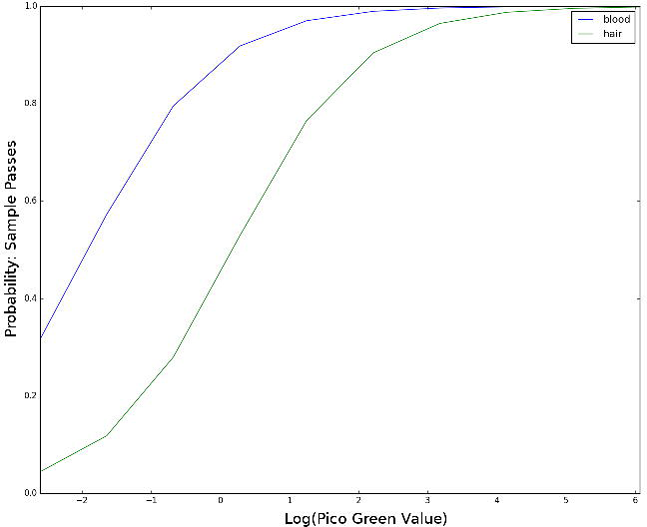
Logistic regression of DNA Sample success. A logistic regression was fit, predicting the probability of a sample passing quality control using DNA concentration (Pico Green) and sample source as independent variables.

